# The diversity of the glycan shield of sarbecoviruses closely related to SARS-CoV-2

**DOI:** 10.1101/2022.08.24.505118

**Authors:** Joel D. Allen, Dylan Ivory, Sophie Ge Song, Wan-ting He, Tazio Capozzola, Peter Yong, Dennis R. Burton, Raiees Andrabi, Max Crispin

## Abstract

The animal reservoirs of sarbecoviruses represent a significant risk of emergent pandemics, as evidenced by the impact of SARS-CoV-2. Vaccines remain successful at limiting severe disease and death, however the continued emergence of SARS-CoV-2 variants, together with the potential for further coronavirus zoonosis, motivates the search for pan-coronavirus vaccines that induce broadly neutralizing antibodies. This necessitates a better understanding of the glycan shields of coronaviruses, which can occlude potential antibody epitopes on spike glycoproteins. Here, we compare the structure of several sarbecovirus glycan shields. Many N-linked glycan attachment sites are shared by all sarbecoviruses, and the processing state of certain sites is highly conserved. However, there are significant differences in the processing state at several glycan sites that surround the receptor binding domain. Our studies reveal similarities and differences in the glycosylation of sarbecoviruses and show how subtle changes in the protein sequence can have pronounced impacts on the glycan shield.

## Introduction

For many years, coronaviruses have been considered a significant threat to public health due to their abundance in animal reservoirs and the severity of disease when zoonosis occurs (Cui et al., 2019). This occurred in 2003 with the SARS-CoV-1 epidemic in Hong Kong (Vijayanand et al., 2004), and in 2010 with a localized outbreak of middle-eastern respiratory syndrome coronavirus (MERS-CoV) (Ramadan and Shaib, 2019). The most severe pandemic resulting from coronavirus zoonosis occurred in 2020 when SARS-CoV-2 spread across the globe, and as of July 2022, has resulted in millions of deaths and half a billion infections worldwide (World Health Organization, 2022). The combined global efforts of researchers worldwide enabled the rapid development vaccines, which have proven to be the most resilient measure in minimizing severe disease and death as lockdowns ease (European Centre for Disease Prevention and Control, 2022; Ssentongo et al., 2022). It is important to note that the development of vaccines in such a short time built on many concepts that were extensively researched prior to the outbreak of COVID-19. Many years of research into RNA as an immunogen delivery mechanism, combined with protein engineering techniques resulted in highly effective vaccines. The COMIRNATY and Spikevax vaccine both employ developments in protein engineering focused around stabilizing the viral spike (S) glycoprotein, maintaining the integrity of neutralizing antibody epitopes (Sanders and Moore, 2021; Scudellari, 2020). In addition to vaccine design, these protein engineering approaches have enabled the in-depth study of the structure and function of the S glycoprotein (Walls et al., 2020; Wrapp et al., 2020).

The coronavirus S protein mediates receptor binding, enabling the virus to enter host cells. Following translation, the S protein consists of a single 200kDa polypeptide chain of over 1200 amino acids, separated into the N-terminal domain (NTD), the receptor binding domain (RBD), fusion peptide (FP), heptad repeat 1 and 2 (HR1/2) and the transmembrane C-terminal domain (Huang et al., 2020). During secretion, the RBD and NTD are separated from the C-terminal elements by proteolytic cleavage, in the case of SARS-CoV-2 this is achieved through the action of the host protease, furin (Bestle et al., 2020). The mature S protein located on the surface of virions consists of a trimer of heterodimers of S1 (containing the NTD and RBD) and S2.

In addition to proteolytic cleavage and maturation, S protein undergoes extensive modified post-translational modifications as the S protein progresses through the ER/Golgi secretory system. The most abundant post-translational modification is N-linked glycosylation, with approximately one third the mass of S protein consisting of N-linked glycans (Walls et al., 2016; Watanabe et al., 2020a). The prevalence of N-linked glycans on viral envelope proteins demonstrates the key roles that N-linked glycans impart upon protein structure and function (Watanabe et al., 2019). Glycans are critical for the correct folding of proteins and stabilize the resultant structure of the viral spike (Varki, 2017). Furthermore, the precise processing state of N-linked glycans is influenced by the surrounding glycan and protein architecture. Thus, the viral genome exerts some control over the processing state (Behrens and Crispin, 2017). While N-linked glycans can contribute to neutralizing antibody epitopes, particularly in HIV (Seabright et al., 2019), their main effect as large, immunologically ‘self’ structures’ is to occlude the underlying protein surface. This means that changes in the glycan shield, with respect to both the position of an N-linked glycan site and the processing state of the attached glycan can modulate viral infectivity and hamper vaccine design efforts (Reis et al., 2021; Vigerust and Shepherd, 2007). Conversely, the presence of under processed glycans on viral glycoprotein immunogens, particularly oligomannosidic forms, can enhance the interaction with the innate immune system and assist trafficking to germinal centers (Tokatlian et al., 2019). Therefore, research into viral biology and vaccine design efforts benefit from an intricate knowledge of the viral glycan shield.

As viral spike proteins are produced by the host, the N-linked glycans attached to mature viral spike glycoproteins will reflect the processing pathway of those cells. Mammalian cells attach N-linked glycans at NxS/T motifs, where x is any amino acid except proline, with this attachment occurring co-translationally, prior to protein folding. The initial stages of mammalian N-linked glycan processing are highly conserved, with the attachment of a pre-assembled glycan containing two N-acetylglucosamine, nine mannose and three glucose monosaccharides. The glucose residues are efficiently cleaved and act as a signal to the calnexin/calreticulin cycle that the protein has folded correctly. Following this, four of the nine mannose residues are removed in the ER and Golgi. From here the pathway diverges, with a multitude of different glycan processing states observed on mature glycoproteins, including the addition of a diverse range of monosaccharides such as fucose and sialic acid (Reily et al., 2019). On the majority of healthy mammalian glycoproteins, the early mannose trimming stages are efficiently performed, and few glycoproteins contain glycans with 5-9 mannose residues. On viral glycoproteins, however, there are a large number of N-linked glycan sites, which results in steric clashes with glycan processing enzymes. Both protein-glycan and protein-protein clashes combine to inhibit N-linked glycan maturation, and oligomannose-type glycans are observed on viral glycoproteins that have exited the secretory system (Watanabe et al., 2019). This is most pronounced on the HIV-1 Envelope glycoprotein (Cao et al., 2017; Struwe et al., 2018); however they have been observed on Influenza HA (Lee et al., 2021), Lassa virus glycoprotein complex (Watanabe et al., 2018), Ebola glycoprotein (Peng et al., 2022), SARS-CoV-1 (Watanabe et al., 2020b), MERS-CoV (Watanabe et al., 2020b) and importantly SARS-CoV-2 (Allen et al., 2021; Brun et al., 2021; Watanabe et al., 2020a; Zhao et al., 2020). The presence of oligomannose-type N-linked glycans on the surface of the spike has been shown to be key indicators of the glycan shield density, and the extent to which the glycan shield occludes immunogenic protein epitopes (Allen et al., 2021).

Differences in the glycan shield can indicate changes in the protein architecture, and therefore a changing antigenic surface. As such it is important to understand the presentation and processing of the N-linked glycans on viral spike glycoproteins. The immunodominant epitope of the SARS-CoV-2 S glycoprotein is the receptor binding domain (RBD) and is poorly shielded by N-linked glycans (Barnes et al., 2020; Cao et al., 2020; Chi et al., 2020; He et al., 2022; Ju et al., 2020; Pinto et al., 2020; Robbiani et al., 2020; Rogers et al., 2020; Seydoux et al., 2020; Shi et al., 2020; Wu et al., 2020; Yuan et al., 2020, 2022). As subsequent variants have demonstrated, this region of the protein is under immune selection pressure, and a few mutations in the RBD can deplete neutralizing antibody binding (Moore and Offit, 2021; Tada et al., 2022). Whilst existing vaccines are effective at preventing serious infection, the titers of neutralizing antibodies from vaccinated individuals are diminished against the variants. These observations are important when considering vaccines that can provide continued protection against an evolving target.

In addition to emerging variants of SARS-CoV-2, it is possible that another zoonotic event involving a new coronavirus will occur in the future. There are many different coronaviruses circulating in nature, many of which share similar sequences to that of SARS-CoV-2 (Letko et al., 2020). Coronaviruses are divided into four genera: alpha, beta, gamma and delta, of which SARS-CoV-2, MERS-CoV and SARS-CoV-1 belong to the betacoronavirus genera. Betacoronavirses can be further classified as a sarbecovirus, merbecovirus, embecovirus or a nobecovirus, with SARS-CoV-1 and SARS-CoV-2 classified as sarbecoviruses. There are many circulating sarbecoviruses, primarily in bats, which possess extensively high sequence similarity to SARS-CoV-2. With a 96% genome identity, RaTG13, found in *Rhinolophus affinis* (bats) in the Yunnan region of China, is the most similar sarbecovirus isolated to that of SARS-CoV-2 (Zhou et al., 2020b). Additionally, a sarbecovirus identified in pangolins, pang17, has a very high similarity (greater than 90%) to SARS-CoV-2 (Lam et al., 2020). The increasing number of isolated sarbecoviruses has resulted in further classification dependent on sequence similarity (**Figure 1A** and **Supplemental Table 2**). SARS-CoV-2, RaTG13 and pang17 are defined into clade 1b whereas SARS-CoV-1 is clade 1a. Other sarbecoviruses circulating in nature also use ACE2 as an entry receptor, including the clade 1a WIV-1 (*Rhinolophus sinicus*) and RsSHC014 (*Rhinolophus sinicus*) (Ge et al., 2013; Zheng et al., 2020). The prevalence of sarbecoviruses in nature that have the potential to spill over into humans warrants the development of a pan-coronavirus vaccine that could be rapidly deployed following an emergent epidemic, to potentially limit the spread of a novel coronavirus.

**Figure 1:**
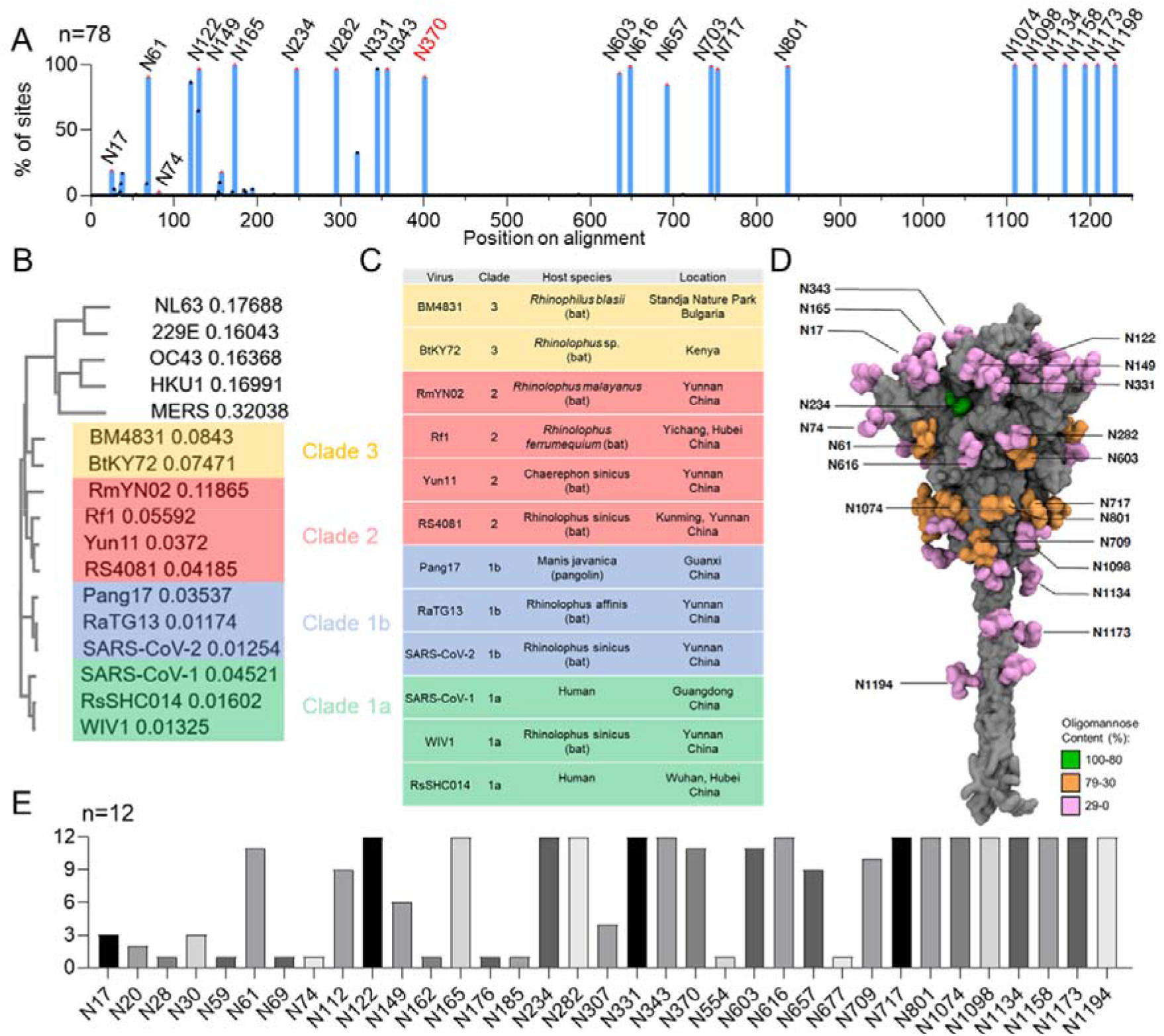
Sarbecoviruses with similar sequences to SARS-CoV-2, and the conservation of N-linked glycosylation sequons across their spike proteins. **A)** Alignment of 78 sarbecovirus S protein sequences. The y-axis represents the proportion of sarbecoviruses that possess an N-linked glycan attachment site, expressed as a percentage of the total sequences used. Peaks corresponding to glycan sites from SARS-CoV-2 are labelled with their position on SARS-CoV-2. N370 is colored red as it is highly conserved, but not present in SARS-CoV-2 **B)** Analysis of the sequence similarity of sarbecoviruses analyzed in this study. Each sarbecovirus is colored according to the clade, which has been classified previously (Cohen et al., 2021). **C)** Table of the sarbecoviruses analyzed in this study, displaying the name, the species it was isolated from and the region in which the isolate was discovered. **D)** Reproduction of previous analysis of the SARS-CoV-2 glycan shield from Allen et al. 2021, determined from the aggregation of data from recombinant protein from multiple sources. The protein is displayed in grey and the glycans are colored according to the abundance of oligomannose-type glycans present at each site (Allen et al., 2021). **E)** Bar chart depicting the number of sarbecoviruses containing an NxS/T motif at a particular site. Each sarbecovirus was aligned to SARS-CoV-2 and the glycan sites are displayed relative to their position on SARS-CoV-2. Linked to Supplemental Table 1

As the N-linked glycans form an integral part of the surface of the spike glycoprotein, it is important that changes in the glycan shield are monitored during the development of potential immunogens that could protect against a broad range of sarbecoviruses. To this end, we selected sarbecoviruses covering multiple clades and introduced mutations that have previously been successfully employed to generate soluble native-like trimers of spike glycoproteins, some of which are used in existing SARS-CoV-2 vaccines. The resultant soluble spike glycoproteins were purified, and the glycosylation analyzed by liquid-chromatography mass spectrometry. By aligning the N-linked glycan sites of the sarbecoviruses to that of SARS-CoV-2, we were able to compare the site-specific glycosylation across the spike glycoproteins. This revealed that the glycosylation, in places, was highly conserved, however other sites are highly variable. To contextualize the changes in glycosylation we generated structural models of the sarbecoviruses and modelled representative glycans onto the structure to investigate the 3-dimensional environment surrounding the N-linked glycan sites. This analysis revealed that the majority of divergent glycosylation patterns occurred on, or proximal to the RBD, such as at N165, suggesting that subtle changes in the amino acid sequence in these regions can have cascading impacts on the glycosylation of the spike protein. Meanwhile, the N-linked glycan sites on the S2 subunit were conserved with respect to both the glycan processing state and the position of the N-linked glycan sequons. These data support observations that the antibodies targeting the S2 region of the protein have the potential to provide a breadth of protection against a range of sarbecoviruses.

## Results

### Comparison of the N-linked glycan positions on sarbecoviruses

The SARS-CoV-2 spike glycoprotein contains 22 N-linked glycosylation sites and is now the most well studied coronavirus spike glycoprotein with regards to glycosylation. The processing of these N-linked glycans is variable (Allen et al., 2021; Casalino et al., 2020), with some sites, such as those towards the C-terminus, processed analogously to host glycoproteins. There are, however, distinct regions of restricted glycan processing, likely resulting from a restrained steric environment, perturbing the ability of glycosidase enzymes to access these sites. A stark example of this is N234, which is enriched for oligomannose-type glycans and we have previously shown that this correlates with a low accessible surface area, as determined using molecular dynamics simulations (Allen et al., 2021). To facilitate comparison with SARS-CoV-2 glycosylation, the sequences of the sarbecovirus spike proteins were aligned to that of SARS-CoV-2, and throughout the manuscript individual sites will be referred to based on their aligned position relative to SARS-CoV-2 (**Supplemental Table 1** and **Supplemental File 1-Sarbecovirus Sequence alignment**)

To compare the presence and location of PNGS across sarbecovirus S proteins, protein sequences for the S protein of 78 sarbecoviruses were obtained from the UniProt database, filtering results for SARS-CoV-2 to investigate sarbecoviruses circulating prior to the outbreak of the COVID-19 pandemic. All S protein sequences were aligned using Clustal Omega. The sequence alignment and list of sarbecovirus S proteins used in this study can be found in **Supplementary File 1: Sarbecovirus Sequence alignment**. In this manner, the conservation of N-linked glycan sites could be compared across the 78 sarbecoviruses **(Figure 1A**). This analysis demonstrated two extremes, either an N-linked glycan site was conserved in the majority of strains analyzed, or highly variable between strains. Key regions of conservation include the aforementioned N234 site, and the two glycan sites located on the SARS-CoV-2 RBD, N331 and N343. Additionally, the N-linked glycan sites on the S2 portion of the protein were conserved on all strains analyzed. Interesting regions of divergence include the N74 glycan site, which was only present on SARS-CoV-2. The N-terminal glycan sites were the most variable with respect to their position, and interestingly some of the emergent SARS-CoV-2 variants have acquired glycan sites in this region which are present on other sarbecoviruses. The gamma variant contains both N17 and N20, as opposed to N17 alone. This N20 site was found in both clade 3 sarbecoviruses used in this study BM4831 and BtKY72, although these strains lack site N17. The gamma variant also contains N188 (Newby et al., 2022), and whilst this site is not present in any of the strains analyzed in this study, 4 sarbecoviruses contained this site in the larger panel. Interestingly, whilst the majority of sarbecoviruses contained N370, SARS-CoV-2 did not. The presence of this site on the RBD likely has profound implications for infectivity and recent molecular dynamics studies have highlighted that the lack of this glycan on SARS-CoV-2 has aided its infectivity (Harbison et al., 2022). Overall, the majority of glycan sites on SARS-CoV-2 are conserved across the sarbecoviruses, hinting at the important role these glycans are playing in maintaining the correct structure and function of the spike glycoprotein.

### Design, expression and purification of sarbecovirus spike glycoproteins

To investigate the variability of the sarbecovirus glycan shield we selected eleven sarbecovirus spike glycoprotein genes and introduced mutations to produce stabilized soluble trimers, using double proline substitutions (2P), a GSAS linker, and a C-terminal trimerization motif. The selected isolates varied in sequence similarity from 70%-97% compared to SARS-CoV-2 at the amino acid level of the spike glycoprotein (**Supplemental Table 2**). In this study, we used sequences corresponding to SARS-CoV-1, WIV1 and RsSHC014 (clade 1a), pang17, RaTG13 and SARS-CoV-2 (clade 1b), RmYN02, Rf1, Yun11 and RS4081 (clade 2) and BM4831 and BtKY72 (clade 3) (**Figure 1B**) (Andersen et al., 2020; Lam et al., 2020; Tao and Tong, 2019; Zhou et al., 2020a). Plasmids encoding the spike glycoproteins were transfected in human embryonic kidney (HEK) 293F cells and the soluble spike glycoproteins were purified from the supernatant using nickel affinity chromatography followed by size exclusion chromatography (SEC). The size exclusion chromatogram displayed a single peak, representing spike glycoprotein trimers.

### Determination of the glycan processing state of sarbecovirus glycan sites

We have previously determined the glycosylation of several coronaviruses, including SARS-CoV-1, MERS-CoV, HKU1 and SARS-CoV-2 (Watanabe et al., 2020a, 2020b) and we employed a similar analytical approach involving in-line liquid chromatography-mass spectrometry (LC-MS). We investigated the highlighted samples in **Figure 1**, however the data for SARS-CoV-2 S protein was obtained from a previous publication (Allen et al., 2021). Three aliquots of the spike glycoproteins were treated separately with trypsin, chymotrypsin, and alpha-lytic protease, with the goal of generating glycopeptides containing a single N-linked glycan site. This enables the glycan processing state of each site to be investigated in a site-specific manner. Following analysis by LC-MS, the compositions of N-linked glycans were determined, and then categorized based on the detected compositions to facilitate comparisons between the different samples. Full glycopeptide identifications for each sample can be found in the **Supplemental file 2: Site-specific glycan analysis**. Compositions corresponding to oligomannose-type glycans are distinct from others as they contain only two N-acetylglucosamine (GlcNAc) residues, whereas complex-type glycans contain at least three. Hybrid-type glycans were defined by the presence of 3-4 HexNAc residues, and 5-6 hexose residues, distinguishing them from complex-type glycans. In this way we identified the proportion of oligomannose-type glycans at each site, for each sarbecovirus (**Figure 2A**). This analysis revealed that although many of the N-linked glycosylation sites are conserved between all sarbecoviruses, the glycan processing of these sites can be highly variable. In addition, there are certain sites that display remarkable conservation across all samples analyzed.

**Figure 2:**
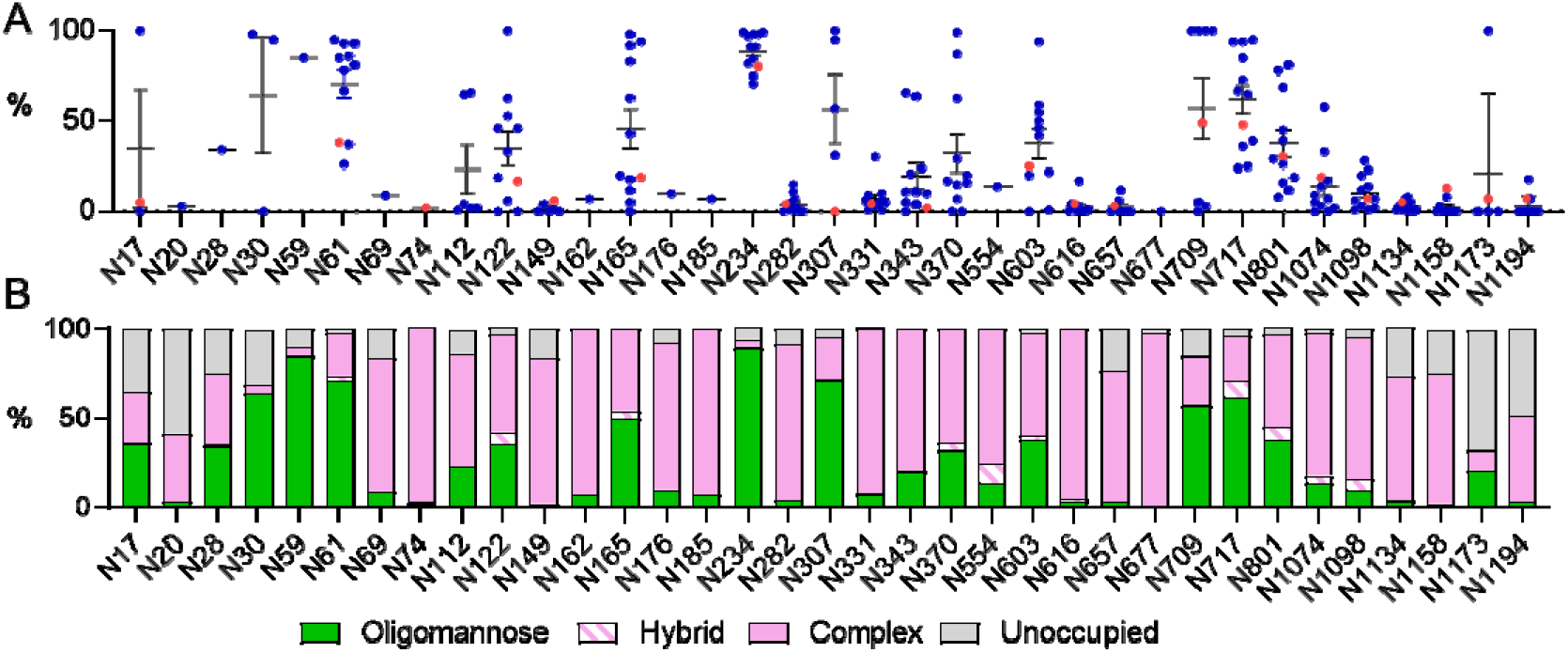
Determination of the site-specific glycosylation of sarbecoviruses by LC-MS. **A)** Sum of the oligomannose-type glycans located at each N-linked glycan sites on the sarbecoviruses analyzed in this study. The sequences for all sarbecoviruses were aligned to SARS-CoV-2 S protein, and the glycan sites are presented aligned to this protein. The oligomannose-type glycan content of previously published site-specific data for SARS-CoV-2 S protein is shown as red dots (Allen et al., 2021). The mean of all strains is displayed as a line and the error bars are +/− SEM, or when only two datasets are present, represent the range of the two datasets. **B)** The averaged glycan processing state of all sarbecoviruses aligned to SARS-CoV-2 S protein. Glycans classified as oligomannose-type consist of compositions containing 2 HexNAc moieties (colored green), hybrid contain 3 or 4 HexNAc and 5 or 6 hexoses, respectively, and is represented as a white bar with pink hatches. Complex-type glycans consist of all remaining detected compositions and are colored pink. The proportion of unoccupied N-linked glycan sites is displayed in grey. Linked to Supplemental Figure 1 and Supplemental Tables 2-5.

The N234 site is located within a sterically restricted environment, proximal to the RBDs at the protomer interface. This glycan has been shown to have important roles in stabilizing the protein fold, controlling RBD dynamics, and removal of this glycan site diminishes the ability of ACE2 to bind (Casalino et al., 2020; Henderson et al., 2020). In all sarbecoviruses analyzed, N234 was occupied by oligomannose-type glycans, ranging from 71% for RS4081, to 99% for BtKY72 (**Figure 2A** and **Supplemental Table 5** and **Supplemental Table 6**). The conservation of the glycan processing provides further evidence of the key role of this glycan in the structure and function of not only SARS-CoV-2, but a broad range of sarbecoviruses. Likewise, the N282 glycan is conserved amongst all sarbecoviruses analyzed but is almost fully occupied by complex-type glycans. The role of this glycan in the structure and function of the S protein is less explored, however the conservation of this site could have important implications due to its proximity to the RBD. Another remarkable region of conservation is the sparsely glycosylated S2 subunit, which ranges from N1074 to N1194. All strains contained the same number and position of N-linked glycosylation sites in the S2 subunit. Additionally, in every sample analyzed the glycan processing was nearly identical, with processed complex-type glycans dominating all sites in this region (**Figure 2A**). The processing of the complex-type glycans were more extensive than on other regions of the glycoprotein, with sites such as N1194 containing elevated levels of glycans consisting of 6 N-acetylhexosamine monosaccharides (**Supplemental Figure 1**). This composition likely corresponds to large tetraantennary glycans and represents extensive glycan processing.

### Establishing a glycan processing consensus reveals trends in sarbecovirus glycosylation

To contextualize the observed differences in the site-specific glycosylation data, we calculated the “consensus” glycosylation across all samples, aligned to the SARS-CoV-2 N-linked glycosylation sites (**Figure 2B** and **Supplemental Figure 1**). Presenting the data in this way enables general trends in glycan processing to be discussed, which can then be compared to outliers within specific strains. This analysis revealed that the glycan processing state of sarbecoviruses is heterogeneous, with oligomannose-type glycans distributed across the spike glycoprotein. This is in contrast to HIV-1 Envelope glycosylation where the processing is more distinct with particular sites on Env consisting entirely of oligomannose-type glycans (Behrens et al., 2017). Across all the samples analyzed, the predominant glycoforms detected was Man_5_GlcNAc_2_ (**Figure 2B, Supplemental Figure 1**). This is consistent with previous analyses of the SARS-CoV-2 spike glycoprotein (Brun et al., 2021; Watanabe et al., 2020a; Zhao et al., 2020). This glycan is an intermediate processing state and is typically present in the cis-Golgi. On the majority of host glycoproteins, this glycan is further processed by the activity of GlcNAc transferase I (GNTI), which then enables the assembly of complex and hybrid-type glycans. This glycan processing bottleneck suggests that the activity of this enzyme is sensitive to the steric environment surrounding the glycan sites, more so than that of the earlier glycan processing enzymes. It has previously been demonstrated that the activity ER-alpha mannosidase I, which converts Man_9_GlcNAc_2_ to Man_8_GlcNAc_2_ can be sterically blocked by proximal glycans and protein, and results in a high abundance of Man_9_GlcNAc_2_ on HIV-1 Env (Pritchard et al., 2015). The glycan shield of sarbecoviruses is less dense than that of HIV-1 Env (Allen et al., 2021; Watanabe et al., 2020b), and likely means that this enzyme is not inhibited to the same extent, but GNTI is instead. This is a key observation for antibody binding, as carbohydrate recognition by antibodies has been shown to favor alpha 1,2 mannose linkages (Scanlan et al., 2002; Williams et al., 2021), which are not present on Man_5_GlcNAc_2_, Additionally, complex-type glycans are found across the protein, with high levels of glycan processing occurring on the RBD glycan sites, N331 and N343. The conserved glycan processing of these glycan sites likely indicates that the glycan shield is sparse around the RBD of all sarbecoviruses analyzed and means that glycan shielding will not impede pan-coronaviruses that use RBD subunits as the immunogenic agents.

Interestingly, populations of N-linked glycan sites were detected that lacked glycan attachment towards the N- and C-terminus of the S proteins. This phenomenon has been reported previously (Allen et al., 2021; Bañó-Polo et al., 2011; Derking et al., 2021) and likely occurs on the C-terminus as a result of the detachment of the translational machinery following translation termination, with the glycosyltransferase remaining attached to the translational machinery. The processing of complex-type glycosylation, with regards to elaboration with additional monosaccharides, such as sialic acid, is driven more by the producer cell used, as the glycosyltransferase expression levels vary from cell to cell, and the complex-type glycan processing present on the sarbecovirus samples is reminiscent of viral glycoproteins previously analyzed from HEK293F cells (Allen et al., 2021). As such, there are limits to the information that can be ascertained from the interpretation of these glycans and would require analysis of virus produced from appropriate cells of origin.

Whilst regions of the glycan shield are highly conserved amongst the majority of sarbecoviruses analyzed, there are several key glycan sites which are highly variable with respect to their glycan processing state, notably N61, N122, N165, N370, N717 and N801. Additional variability was observed at sites such as N17, N30 and N307 however, these sites were less conserved across sarbecoviruses analyzed. The variation in glycan processing at these sites varied from entirely oligomannose-type to entirely complex-type (**Supplemental Tables 3 to Supplemental Table 6)**. For example, Pang17, contains 98% oligomannose-type glycans at N165, whereas RaTG13 contains 5% at the same site (**Figure 2A**). The N165 glycan has been shown to have an important role in mediating the conformation of the RBD, facilitating the RBD-up position, which is favorable for receptor binding, and also exposes neutralizing antibody epitopes. As such, changes in the processing of this glycan may be indicative of differential RBD dynamics between the sarbecoviruses (Casalino et al., 2020; Chawla et al., 2022).

### The extent of clade-specific glycan processing of sarbecoviruses

As there are regions of the glycan shield of sarbecoviruses that are extremely variable, we sought to investigate whether closely linked sarbecoviruses have convergent glycosylation, and whether the variability arises between clades. Using the classification outlined in **Figure 1** we compared the site-specific glycosylation between sarbecoviruses in clade 1a, clade 1b, clade 2 and clade 3 (**Figure 3**). Clades 1 and 1b contained the highest proportion of oligomannose-type glycans (**Figure 3A** and **B**), and clade 3 the least (**Figure 3D**). This can be seen most prominently on glycan sites located towards the C-terminus of the S1 domain, such as N717 and N801. In clade 1a, these sites are almost fully occupied by oligomannose-type glycans, for example on the clade 1a RsSCHC014 the N717 site contains 95% oligomannose-type glycans whereas on BtkY72, of clade 3, this same site is only occupied by oligomannose-type glycans on 26% of sites (**Supplemental Table 3** and **6**). Sites such as N717 and N801 have been shown to form the epitopes of glycan binding antibodies that target oligomannose-type glycans (Williams et al., 2021). These data suggests that these glycan epitopes may not be conserved across sarbecoviruses, and may not provide broad protection, although the antibodies may still bind at other regions of the trimer.

**Figure 3:**
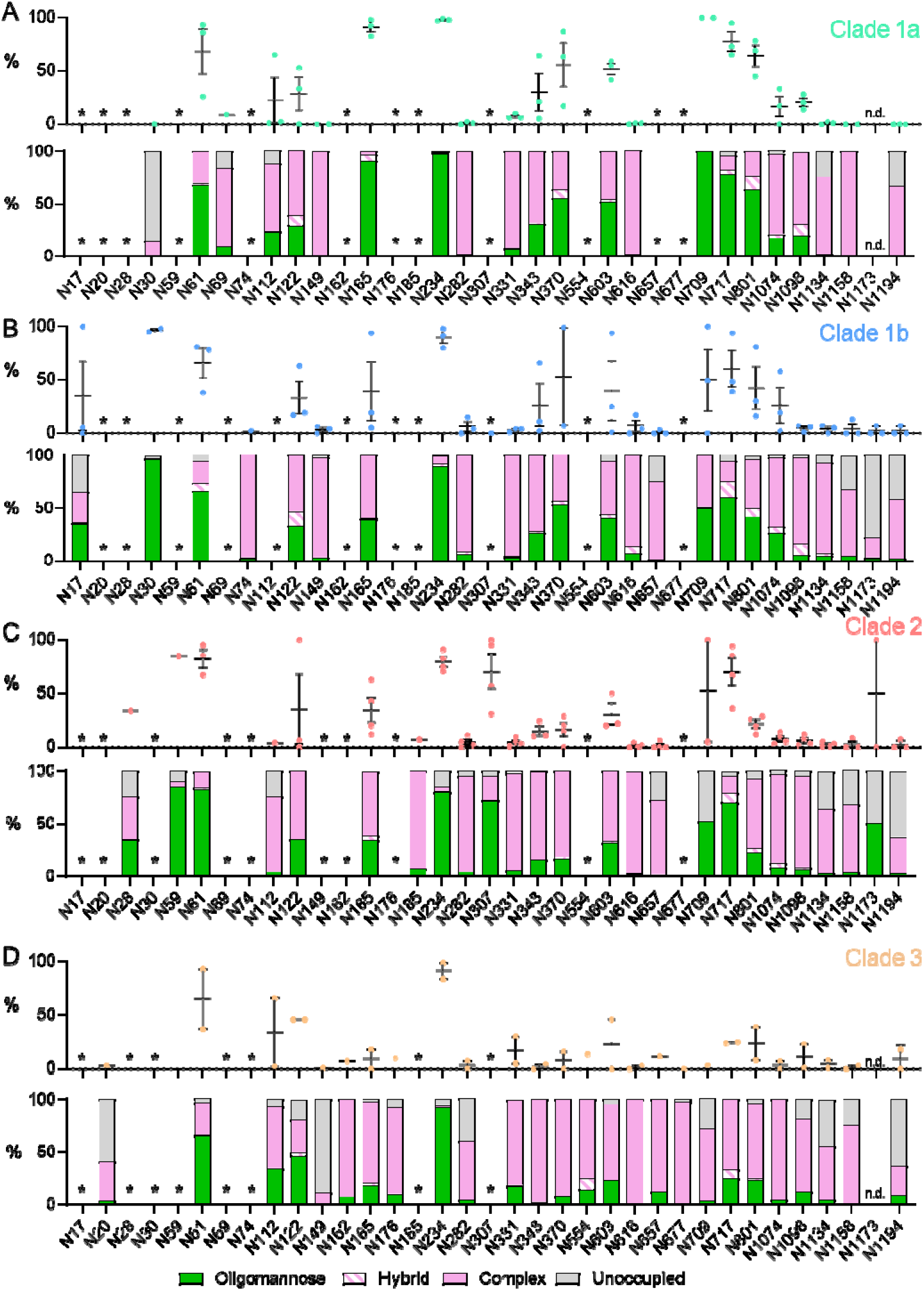
Clade-specific glycan processing of sarbecoviruses. **A)** Site-specific glycosylation of clade 1a sarbecoviruses, with the data displayed in an identical manner to Figure 2, with the symbols representing the oligomannose-type glycan content of individual strains, and the bar graph representing the consensus glycosylation pattern at each site. **B)** Site-specific glycosylation of clade 1b sarbecoviruses. **C)** Site-specific glycosylation of clade 2 sarbecoviruses. **D)** Site-specific glycosylation of clade 3 sarbecoviruses. Sites that are not present in a particular clade are labelled with an asterisk. Sites where the site-specific glycosylation could not be determined. Linked to Supplemental Tables 2, 3, 4 and 5.

In addition to glycan sites towards the base of the trimer, many of the N-linked glycan sites displaying variable processing states are located around the RBD. The N165 site has been shown to be a sensitive reporter of RBD dynamics (Chawla et al., 2022), and is highly processed in clade 3, whereas clade 1a is almost entirely populated by oligomannose-type glycans. These same sites vary between individual strains in clade 1b and clade 2 and as such the glycan processing state of this site is not clade specific. The high variability of glycan processing, despite a broad conservation in the position of N-linked glycan sites suggests that glycan position alone is not a predictor of glycan processing state. It is therefore important to understand the presentation of the glycan in its 3-dimensional environment to understand how the glycan shield can vary between different strains which are broadly conserved at the amino acid level.

### Generating molecular models of sarbecovirus glycan shields

As the glycan shield varies in composition despite a broad conservation of N-linked glycan sequon position, we sought to contextualize the site-specific glycosylation mapping the glycan shield onto the underlying protein structure (**Figure 4**). To enable this, a combination of molecular modelling tools were used. First, the sequences of the sarbecovirus spike proteins used for the generation of the recombinant protein were uploaded to SWISS-MODEL (Bienert et al., 2017; Studer et al., 2020, 2021; Waterhouse et al., 2018).The SWISS-MODEL template library (SMTL version 2022-04-27, PDB release 2022-04-22) was searched with BLAST (Camacho et al., 2009) and HHblits (Steinegger et al., 2019) for evolutionary related structures matching the target sequence.

**Figure 4:**
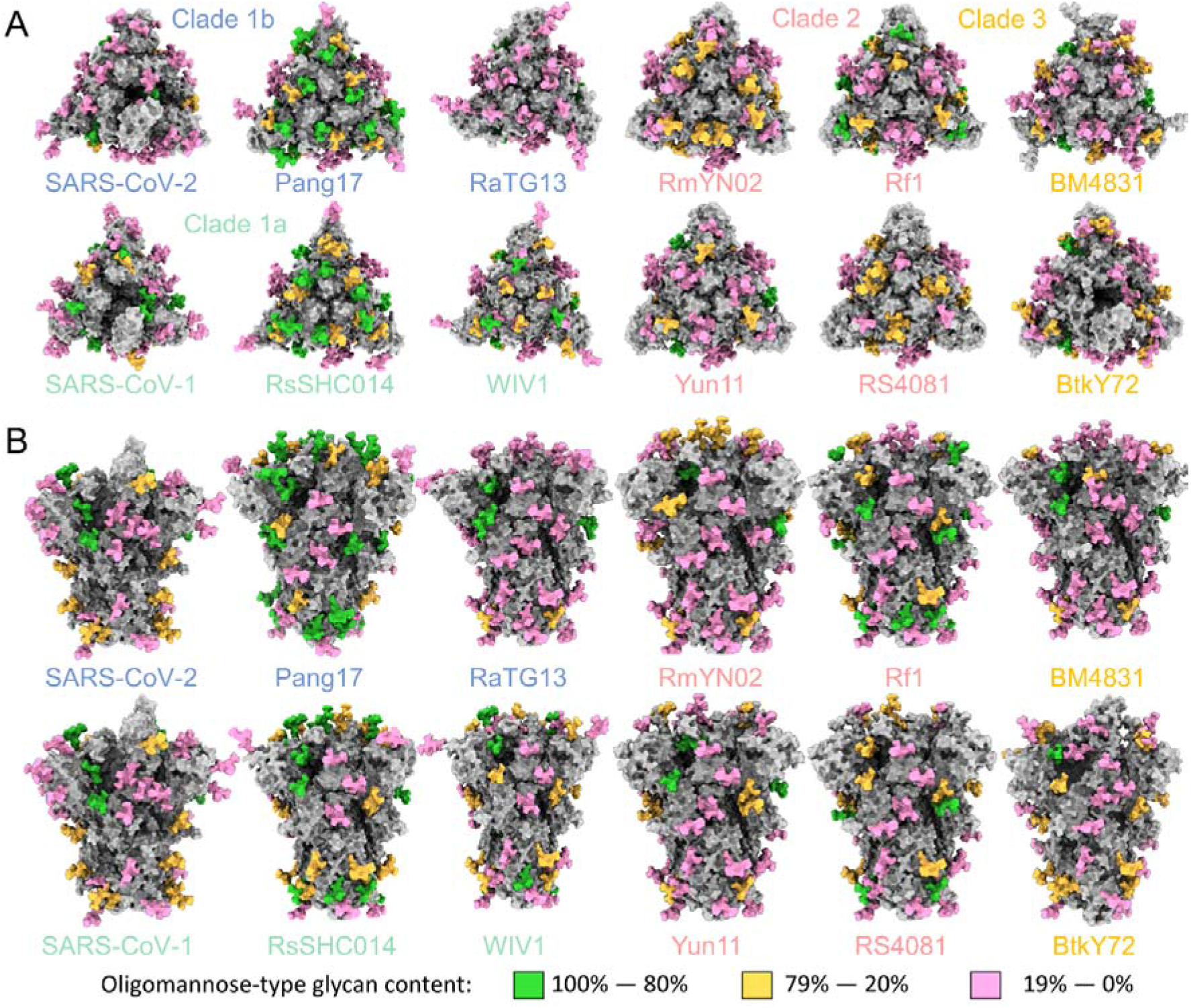
Modelling the glycan shield of sarbecoviruses with their site-specific glycosylation. 3D maps of the sarbecoviruses glycan shields are displayed top down (A) and side on (B). All models were constructed using Swiss model, GlycoShield and the mass spectrometry data displayed in Figure 3. Each model displays the protein sequence as grey. A representative Man_5_GlcNAc_2_ glycan was mapped onto each potential N-linked glycosylation site and is colored according to the oligomannose-type glycan content at each site, with 80% and above colored green, between 79% and 20% colored orange, and below 20% colored pink. The C-terminal region of the Spike protein was not resolved in the templates used to generate the models, and so is not included.

These templates do not contain glycans and so we used an additional tool to attach a representative N-linked glycans at each site. As Man_5_GlcNAc_2_ is the most abundant single composition on all samples, we modelled this glycan on every site using GlycoSHIELD (Gecht et al., 2022). This approach enabled 3D maps of the glycan shield to be generated for the 11 sarbecoviruses analyzed in this manuscript, as well as for SARS-CoV-2, analyzed previously (Allen et al., 2021). As many of the templates used to generate these maps did not contain a portion of the C-terminal domain, this was not included in our models, and the three C-terminal glycan sites are not included. As these are processed in a similar manner across all samples analyzed, and they consist of almost exclusively complex-type glycans, the processing of these sites is likely not influenced by glycan or protein clashes. Qualitatively, these models demonstrate the variability in the glycan shield that was shown with the site-specific analysis. Within clades, the glycan processing is variable. Two examples of this include SARS-CoV-2 and the closely related pangolin coronavirus, pang17, from clade 1b, as well as the clade 2 Rf1 sarbecovirus, compared to the other strains in this clade. It is important to note that as these models were generated based on previously resolved templates and do not represent experimentally determined structures. Fully glycosylated models of the sarbecoviruses can be found at 10.5281/zenodo.7015311.

### Variable glycan processing, despite a conservation of N-linked glycan site positions

From the 3D glycosylation maps generated, the starkest differences in glycosylation were observed in clade 1b. This clade includes SARS-CoV-2, RaTG13 and pang17. Of all the sarbecoviruses analyzed, RaTG13 and pang17 share the most amino acids with SARS-CoV-2 and understanding how the glycosylation diverges in such similar viruses is a key part of exploring the antigenic diversity of the sarbecovirus glycan shield. To compare these viruses, the glycosylation maps corresponding to clade 1b sarbecoviruses is displayed in **Figure 5**. At the majority of sites, RaTG13 and SARS-CoV-2 are similarly glycosylated, however there is an elevated level of oligomannose-type glycans across the pang17 spike (**Figure 5B**). Whilst SARS-CoV-2 contains distinct glycosylation positions, lacking N370 and containing N74, RaTG13 and pang17 contain the exact same number and position of N-linked glycosylation sites.

**Figure 5:**
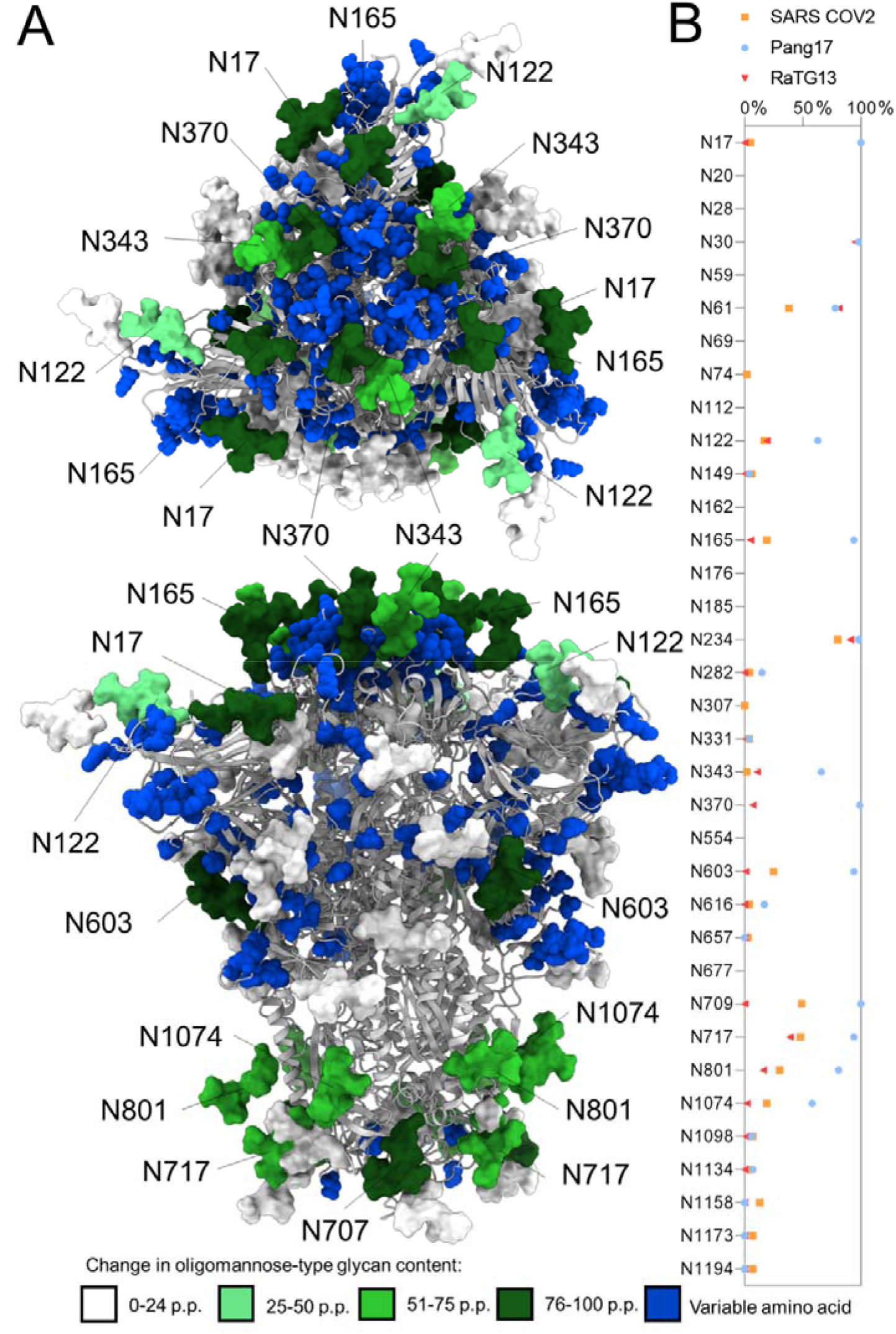
Glycan shield map of RaTG13-CoV to investigate the distinct glycan processing observed in clade 1b sarbecoviruses. **A)** Reproduction of the RaTG13 model generated in Figure 4, with the glycans recolored according to the percentage point difference in oligomannose-type glycans between RaTG13, and pang17, with a positive number representing an increase in the oligomannose-type glycan content of pang17 relative to RaTG13. The protein sequence is displayed as a cartoon depiction, with discrepancies in the amino acid sequence between RaTG13 and pang17 represented as blue spheres. Sites displaying increased oligomannose-type glycans on pang17 are labelled, with the top panel representing a top-down view, and the bottom panel a side on view. **B)** Comparing the site-specific oligomannose-type glycan content of clade 1b sarbecoviruses, SARS-CoV-2, pang17 and RaTG13.

To understand how the observed variability in glycan processing could be arising, we utilized the 3D glycosylation maps generated in **Figure 4**. In addition to the N-linked glycosylation sites, we compared regions of the protein sequence that differed between RaTG13 and pang17 (**Figure 5A**). Across the pang17 spike glycoprotein, there were several sites that showed a significant increase in the abundance of oligomannose-type glycans compared to both SARS-CoV-2 and RaTG13: N17, N61, N122, N165, N343, N370, N603, N709, N717, N801 and N1074 (**Figure 5B**). Other regions of the glycan shield are conserved, such as the presentation of oligomannose-type glycans at N234, and more processed regions at N282 and the C-terminal sites. Highlighted in blue in **Figure 5** are amino acids that differ between RaTG13 and pang17. Whilst the majority of amino acids are conserved, with 93.19% sequence identity (**Supplemental table 2**), there are clusters of variable amino acids across the spike. The most variable region is located on and around the RBD domain, in a similar manner to the accumulation of mutations on the emergent SARS-CoV-2 variants. This region around the apex of the trimer (**Figure 5A, top panel**) is also where the N-linked glycosylation sites are the most variable, with respect to glycans processing. Several of these sites located in and around the RBD display an extensive restriction in glycan processing, most notably N165, N370 and N343. Additionally, the N603 glycan site displays a similar increase in oligomannose-type glycans and the amino acids around this region are variable as well. There are several sites towards the C-terminus display elevated oligomannose-type glycans however, the protein sequence in this region is less variable. The increase in oligomannose-type glycans at these sites is not as pronounced as those around the apex, and RaTG13 and SARS-CoV-2 both contain oligomannose-type glycans at these sites. These results demonstrate that despite a broad conservation of amino acids across the clade 1b sarbecoviruses, a limited number of mutations in key regions of the spike are impacting the glycan shield. As changing levels of oligomannose-type glycans can act as reporters for changes in the protein architecture (Allen et al., 2021; Behrens et al., 2017), these results suggest that changes in the amino acid structures that modulate the structure of the protein will have impacts upon the glycan processing across the glycan shield. Sites such as N165 have been shown to be sensitive to changes in the protein sequence, for example, the introduction of additional stabilizing mutations into the Wuhan hu1 SARS-CoV-2 spike, termed the HexaPro construct, demonstrated similar changes in glycosylation at this site (Chawla et al., 2022; Hsieh et al., 2020).

### Perspectives

The propensity of SARS-CoV-2 to mutate and generate new variants of concern highlights the importance of investigating the molecular architecture of similar sarbecoviruses, to prepare for a potential species crossover in the future. The sarbecoviruses investigated in this study share similar sequences with SARS-CoV-2, and are being investigated for use in formats for pan-sarbecovirus vaccines (Pinto et al., 2020). The goal of this study was to investigate the variability in the glycan shield of sarbecoviruses, as they constitute one third of the mass of the surface of the spike glycoprotein, and alterations in their presence and processing will alter the antigenic surface of the viral spike. With regards to the position of potential N-linked glycosylation sites, the majority of sites were conserved with SARS-CoV-2. The N-terminal region displayed the most variability, and this is reflected in SARS-CoV-2 variants, with the gamma variant containing the N20 glycosylation site not found in the original Wuhan strain. Analysis of the glycan processing of the sarbecoviruses revealed regions of both conserved and divergent glycan processing. The N234 site likely plays a key role in the stability and function of the spike protein (Casalino et al., 2020), and its position and processing state were conserved across all samples analyzed. The most remarkable conservation was observed in the S2 region of the protein, with sites N1074, N1098, N1134, N1158, N1173 and N1198 conserved across all samples with respect to both position and processing state, exhibiting low levels of oligomannose-type glycans. Conversely, some regions display highly divergent glycan processing, with the conserved N165 glycan site displaying extensive heterogeneity. We generated 3D maps of the sarbecovirus glycan shields to contextualize the changes in glycosylation, and for three highly similar sarbecoviruses; SARS-CoV-2, RaTG13 and pang17, we showed that slight modifications in the amino acid sequence can result in distinct glycosylation profiles, most notably on and around the RBD.

Our observations provide insight into regions that may prove more promising in the design of pan-sarbecovirus vaccines, demonstrating that subtle changes in the amino acids of the RBD can have profound impacts upon the architecture of the spike, which impacts the processing of N-linked glycans. This is also seen in the continued evolution of SARS-CoV-2, where mutations in the spike protein are focused on the RBD and S1 domains. This is in response to these regions being the immunodominant regions of the spike glycoprotein, and subtle changes in this region can diminish the ability of neutralizing antibodies to recognize new variants. Conversely the S2 region is conserved and may provide a more attractive target for vaccine design. Indeed, several studies have highlighted the potential for broad coronavirus antibody recognition and neutralization by exploiting this domain (Hurlburt et al., 2022; Jette et al., 2021; Lv et al., 2020; Shah et al., 2021; Wang et al., 2021). It is important to note that these antibodies are not as potent as RBD specific neutralizing antibodies. The discovery of many coronaviruses in animal reservoirs suggests that, in a similar manner to influenza, coronavirus induced pandemics are of considerable likelihood in the future and understanding the antigenic surface of these viruses and how it can change is important to consider when preparing for future outbreaks.

## Supporting information

Supplemental File 1- Sarbecovirus Sequence alignment

Supplemental File 2- Site specific glycan analysis

## Acknowledgments

This work was supported by the International AIDS Vaccine Initiative (IAVI) through grant INV-008352/OPP1153692 funded by the Bill and Melinda Gates Foundation (M.C.). We also gratefully acknowledge support from the University of Southampton Coronavirus Response Fund (M.C.), a donation from the Bright Future Trust (M.C). NIH NIAID CHAVD (UM1 AI44462 to D.R.B.), IAVI Neutralizing Antibody Center, the Bill and Melinda Gates Foundation (OPP 1170236 and INV-004923 to D.R.B). This work was also supported by the John and Mary Tu Foundation and the James B. Pendleton Charitable Trust (D.R.B.).

## Author contributions

Conceptualization: **JDA, MC.** Formal analysis: **JDA, DI.** Investigation **JDA DI SGS WH TC.** Resources: **SGS, WH, TC, PY, DRB, RA**. Data curation: **JDA**. Writing- original draft **JDA.** Funding acquisition **MC RA DRB.** All authors contributed to reviewing and editing the manuscript.

## STAR methods

### Data

Raw files, protein sequences and glycan library used for the site-specific glycan analysis are deposited on the MASSive server under the following ID: ftp://massive.ucsd.edu/MSV000090155/

Glycosylated models of the coronaviruses used in this study are deposited: 10.5281/zenodo.7015311

The spike constructs include SARS-CoV-2 (residues 1–1208; GenBank ID: MN908947), SARS-CoV-1 (residues 1–1190; GenBank ID: AAP13567), RaTG13 (residues 1-1204, GenBank ID: QHR63300.2), Pang17 (residues 1-1202, GenBank ID: QIA48632.1), WIV1 (residues 1-1191, GenBank ID: KF367457), RsSHC014 (residues 1-1191, GenBank ID: AGZ48806.1), BM48-31 (residues 1-1194, GenBank ID: NC_014470.1), BtKY72 (residues 1-1193, GenBank ID: KY352407), RmYN02 (residues 1-1165, GISAID ID: EPI_ISL_412977), Rf1 (residues 1-1176, GenBank ID: DQ412042.1), Rs4081 (residues 1-1176, GenBank ID: KY417143.1), Yun11 (residues 1-1176, GenBank ID: JX993988).

Any additional information required to reanalyze the data reported in this paper is available from the lead contact upon request.

## METHOD DETAILS

### Potential N-linked glycan conservation and alignment search

To investigate the distribution of potential N-linked glycan sites on sarbecoviruses, the UniProt database was searched with two terms: “sarbecovirus” “spike” -“severe acute respiratory syndrome coronavirus 2 (2019-nCoV) (SARS-CoV-2)” “bat” and “sarbecovirus” “spike” -"severe acute respiratory syndrome coronavirus 2 (2019-nCoV) (SARS-CoV-2)” “pangolin”. This approach was taken to limit the results to sarbecoviruses in animals and not SARS-CoV-2 circulating in animals. A total of 78 sequences were obtained and were aligned using Clustal Omega. The aligned sequences were then searched for PNGS, and the percentage of sites was determined. The full list of sequences is available in **Supplemental File 1-Sarbecovirus Sequence alignment**.

### Protein production and DNA template design

The expression plasmids of soluble spike ectodomain proteins were constructed by DNA fragments synthesized at GeneArt (Thermo Fisher Scientific) followed by cloning into the phCMV3 vector by Gibson assembly. The soluble spike proteins were stabilized in the trimeric prefusion state by introducing double proline substitutions (2P) in the S2 subunit, replacing the furin cleavage sites by a GSAS linker, as well as incorporating the trimerization motif T4 fibritin at the C terminus of the spike proteins. The HRV-3C protease cleavage site, 6×His-Tag and AviTag spaced by GS linkers were added to the C terminus for protein purification and biotinylation.

For protein expression, 350ug of the plasmids encoding spikes were transfected into 1L HEK-293F cells at 1 million cells/ml using Transfectagro (Corning) and 40K polyethylenimine (PEI) (1 mg/ml). The plasmid and transfection reagents were combined and filtered before PEI was added. The mixture solution was incubated at room temperature for 20-30 min before being added into cells. After 4 days, the supernatant was centrifuged and filtered, followed by loading onto columns with HisPur Ni-NTA resin (Thermo Fisher Scientific). The resin-bound protein was washed (25 mM imidazole, pH 7.4) and eluted using 25 ml elution buffer (250 mM imidazole, pH 7.4). The eluate was buffer-exchanged into PBS and further purified through size-exclusion chromatography (SEC) by Superdex 200 (GE Healthcare).

### Site-specific glycan analysis by LC-MS

Three aliquots of sarbecovirus were denatured for 1h in 50 mM Tris/HCl, pH 8.0 containing 6 M of urea and 5 mM dithiothreitol (DTT). The denatured proteins were alkylated by adding 20 mM iodoacetamide (IAA) and incubated for 1h in the dark, followed by a 1h incubation with 20 mM DTT to eliminate residual IAA. The alkylated Env proteins were buffer-exchanged into 50 mM Tris/HCl, pH 8.0 using Vivaspin columns (3 kDa) and two of the aliquots were digested separately overnight using trypsin, chymotrypsin (Mass Spectrometry Grade, Promega) or alpha lytic protease (Sigma Aldrich) at a ratio of 1:30 (w/w). The next day, the peptides were dried and extracted using C18 Zip-tip (MerckMilipore). The peptides were dried again, re-suspended in 0.1% formic acid and analyzed by nanoLC-ESI MS with an Ultimate 3000 HPLC (Thermo Fisher Scientific) system coupled to an Orbitrap Eclipse mass spectrometer (Thermo Fisher Scientific) using stepped higher energy collision-induced dissociation (HCD) fragmentation. Peptides were separated using an EasySpray PepMap RSLC C18 column (75 μm × 75 cm). A trapping column (PepMap 100 C18 3μM x 75μM x 2cm) was used in line with the LC prior to separation with the analytical column. The LC conditions were as follows: 280-minute linear gradient consisting of 4-32% acetonitrile in 0.1% formic acid over 260 minutes followed by 20 minutes of alternating 76% acetonitrile in 0.1% formic acid and 4% Acn in 0.1% formic acid, used to ensure all the sample had eluted from the column. The flow rate was set to 300 nL/min. The spray voltage was set to 2.5 kV and the temperature of the heated capillary was set to 40 °C. The ion transfer tube temperature was set to 275 °C. The scan range was 375−1500 m/z. Stepped HCD collision energy was set to 15, 25 and 45% and the MS2 for each energy was combined. Precursor and fragment detection were performed using an Orbitrap at a resolution MS1= 120,000. MS2= 30,000. The AGC target for MS1 was set to standard and injection time set to auto which involves the system setting the two parameters to maximize sensitivity while maintaining cycle time. Full LC and MS methodology can be extracted from the appropriate Raw file using XCalibur FreeStyle software or upon request.

Glycopeptide fragmentation data were extracted from the raw file using Byos (Version 3.5; Protein Metrics Inc.). The glycopeptide fragmentation data were evaluated manually for each glycopeptide; the peptide was scored as true-positive when the correct b and y fragment ions were observed along with oxonium ions corresponding to the glycan identified. The MS data was searched using the Protein Metrics 305 N-glycan library with sulfated glycans added manually. The relative amounts of each glycan at each site as well as the unoccupied proportion were determined by comparing the extracted chromatographic areas for different glycotypes with an identical peptide sequence. All charge states for a single glycopeptide were summed. The precursor mass tolerance was set at 4 ppm and 10 ppm for fragments. A 1% false discovery rate (FDR) was applied. The relative amounts of each glycan at each site as well as the unoccupied proportion were determined by comparing the extracted ion chromatographic areas for different glycopeptides with an identical peptide sequence. Glycans were categorized according to the composition detected.

HexNAc(2)Hex(10+) was defined as M9Glc, HexNAc(2)Hex(9−5) was classified as M9 to M3. Any of these structures containing a fucose were categorized as FM (fucosylated mannose). HexNAc(3)Hex(5−6)X was classified as Hybrid with HexNAc(3)Hex(5-6)Fuc(1)X classified as Fhybrid. Complex-type glycans were classified according to the number of HexNAc subunits and the presence or absence of fucosylation. As this fragmentation method does not provide linkage information compositional isomers are grouped, so for example a triantennary glycan contains HexNAc 5 but so does a biantennary glycans with a bisect. Core glycans refer to truncated structures smaller than M3. M9glc- M4 were classified as oligomannose-type glycans.

### Model generation: Template Search

Template search with BLAST and HHblits was performed against the SWISS-MODEL template library (SMTL, last update: 2022-04-27, last included PDB release: 2022-04-22). The target sequence was searched with BLAST against the primary amino acid sequence contained in the SMTL. An initial HHblits profile was built using the procedure outlined in (Steinegger et al., 2019), followed by 1 iteration of HHblits against Uniclust30 (Mirdita et al., 2017). The obtained profile was then searched against all profiles of the SMTL.

### Model generation: Model Building

Models are built based on the target-template alignment using ProMod3 (Studer et al., 2021). Coordinates which are conserved between the target and the template are copied from the template to the model. Insertions and deletions are remodeled using a fragment library. Side chains are then rebuilt. Finally, the geometry of the resulting model is regularized by using a force field. The global and per-residue model quality has been assessed using the QMEAN scoring function (Studer et al., 2020).The quaternary structure annotation of the template is used to model the target sequence in its oligomeric form. The method (Bertoni et al., 2017) is based on a supervised machine learning algorithm, Support Vector Machines (SVM), which combines interface conservation, structural clustering, and other template features to provide a quaternary structure quality estimate (QSQE). To map the N-linked glycans to the sarbecovirus templates GlycoSHIELD was used to graft glycan conformers derived from extensive molecular dynamics simulations (Gecht et al., 2022). A representative N-linked glycan was used Man_5_GlcNac_2_. The grafting procedure was performed using a cutoff radius of 0.7 Å.

