## Supplemental Information for "The diversity of the glycan shield of sarbecoviruses closely related to SARS-CoV-2"

**Supplemental Table 1: Conservation of SARS-CoV-2 PNGS across 78 sarbecovirus sequences**

| Position on alignment | SARS-CoV-2 glycan position | % of sites |
| --- | --- | --- |
| 25 | 17 | 19% |
| 69 | 61 | 91% |
| 82 | 74 | 3% |
| 130 | 122 | 97% |
| 157 | 146 | 18% |
| 173 | 165 | 100% |
| 247 | 234 | 97% |
| 295 | 282 | 97% |
| 344 | 331 | 97% |
| 356 | 343 | 97% |
| 401 | 370 | 91% |
| 635 | 603 | 94% |
| 648 | 616 | 99% |
| 692 | 657 | 85% |
| 745 | 709 | 99% |
| 753 | 717 | 97% |
| 837 | 801 | 99% |
| 1110 | 1074 | 100% |
| 1134 | 1098 | 100% |
| 1170 | 1134 | 100% |
| 1194 | 1158 | 100% |
| 1209 | 1173 | 100% |
| 1230 | 1194 | 100% |

**Supplemental Table 2: Coronavirus sequence similarity displayed as a percentage identity matrix**

|  | NL63 | 229E | OC43 | HKU1 | MERS | BM4831 | BtKY72 | Pang17 | RaTG13 | SARS-CoV-2 | RmYN02 | SARS-CoV-1 | RsSHC014 | WIV1 | Rf1 | Yun11 | RS4081 |
| --- | --- | --- | --- | --- | --- | --- | --- | --- | --- | --- | --- | --- | --- | --- | --- | --- | --- |
| NL63 | 100 | 66 | 31 | 31 | 29 | 31 | 31 | 29 | 30 | 30 | 30 | 30 | 30 | 30 | 31 | 30 | 31 |
| 229E | 66 | 100 | 32 | 33 | 32 | 31 | 32 | 31 | 32 | 32 | 32 | 32 | 32 | 32 | 32 | 32 | 32 |
| OC43 | 31 | 32 | 100 | 67 | 37 | 36 | 35 | 35 | 35 | 35 | 36 | 35 | 36 | 36 | 37 | 36 | 36 |
| HKU1 | 31 | 33 | 67 | 100 | 36 | 35 | 34 | 34 | 35 | 35 | 36 | 35 | 35 | 36 | 36 | 34 | 35 |
| MERS | 29 | 32 | 37 | 36 | 100 | 35 | 35 | 34 | 35 | 35 | 35 | 35 | 35 | 35 | 35 | 35 | 35 |
| BM4831 | 31 | 31 | 36 | 35 | 35 | 100 | 84 | 73 | 73 | 73 | 72 | 76 | 76 | 76 | 75 | 76 | 76 |
| BtKY72 | 31 | 32 | 35 | 34 | 35 | 84 | 100 | 74 | 74 | 74 | 73 | 77 | 77 | 77 | 77 | 78 | 77 |
| Pang17 | 29 | 31 | 35 | 34 | 34 | 73 | 74 | 100 | 93 | 93 | 76 | 78 | 78 | 78 | 77 | 77 | 77 |
| RaTG13 | 30 | 32 | 35 | 35 | 35 | 73 | 74 | 93 | 100 | 98 | 76 | 78 | 78 | 78 | 77 | 77 | 77 |
| SARS2 | 30 | 32 | 35 | 35 | 35 | 73 | 74 | 93 | 98 | 100 | 76 | 77 | 78 | 78 | 77 | 77 | 77 |
| RmYN02 | 30 | 32 | 36 | 36 | 35 | 72 | 73 | 76 | 76 | 76 | 100 | 74 | 74 | 74 | 79 | 79 | 79 |
| SARS1 | 30 | 32 | 35 | 35 | 35 | 76 | 77 | 78 | 78 | 77 | 74 | 100 | 90 | 92 | 79 | 81 | 81 |
| RsSHC014 | 30 | 32 | 36 | 35 | 35 | 76 | 77 | 78 | 78 | 78 | 74 | 90 | 100 | 97 | 80 | 81 | 81 |
| WIV1 | 30 | 32 | 36 | 36 | 35 | 76 | 77 | 78 | 78 | 78 | 74 | 92 | 97 | 100 | 80 | 81 | 81 |
| Rf1 | 31 | 32 | 37 | 36 | 35 | 75 | 77 | 77 | 77 | 77 | 79 | 79 | 80 | 80 | 100 | 90 | 89 |
| Yun11 | 30 | 32 | 36 | 34 | 35 | 76 | 78 | 77 | 77 | 77 | 79 | 81 | 81 | 81 | 90 | 100 | 92 |
| RS4081 | 31 | 32 | 36 | 35 | 35 | 76 | 77 | 77 | 77 | 77 | 79 | 81 | 81 | 81 | 89 | 92 | 100 |

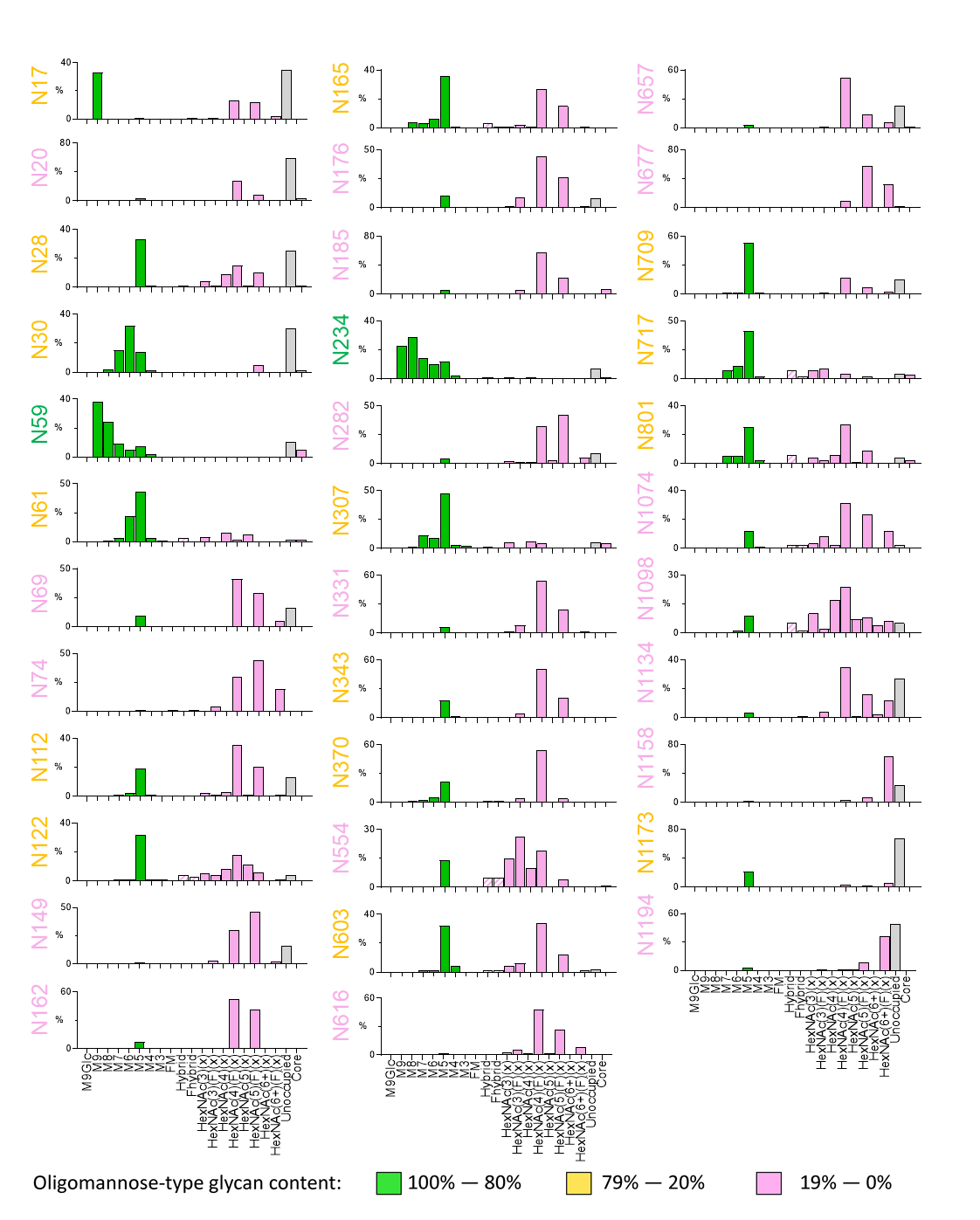
**Supplemental Figure 1: Consensus site-specific glycan analysis of sarbecovirus glycan shields.** The consensus site-specific glycosylation of all sarbecoviruses analyzed in this study. Each sarbecovirus was aligned to SARS-CoV-2 and where a conserved glycan site was present, the detected compositions were averaged. The compositions detected by LC-MS were grouped, as outlined in the materials and methods. Oligomannose-type glycans are displayed as green, hybrid-type glycans are displayed as white, with pink cross hatching and complex-type glycans displayed as pink bars. The proportion of unoccupied N-linked glycan sites is displayed as a grey bar. Each site label is colored according to the oligomannose-type glycan content.

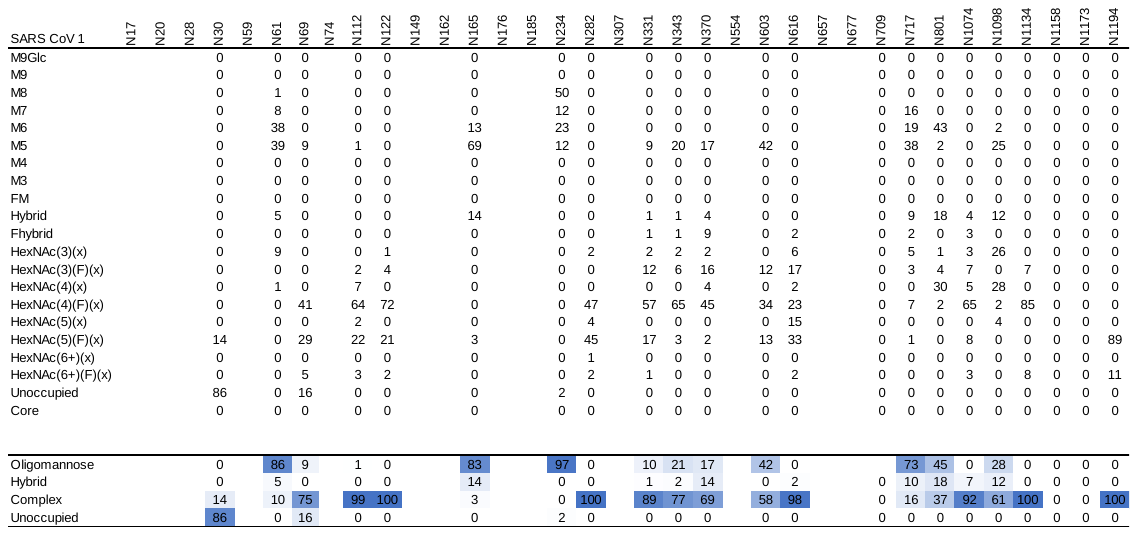

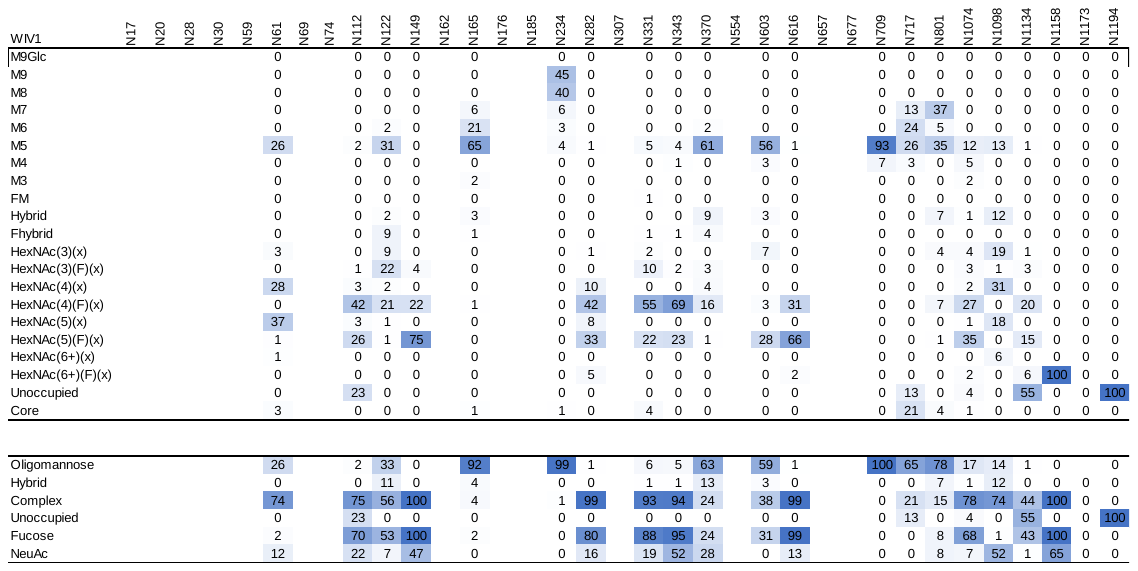

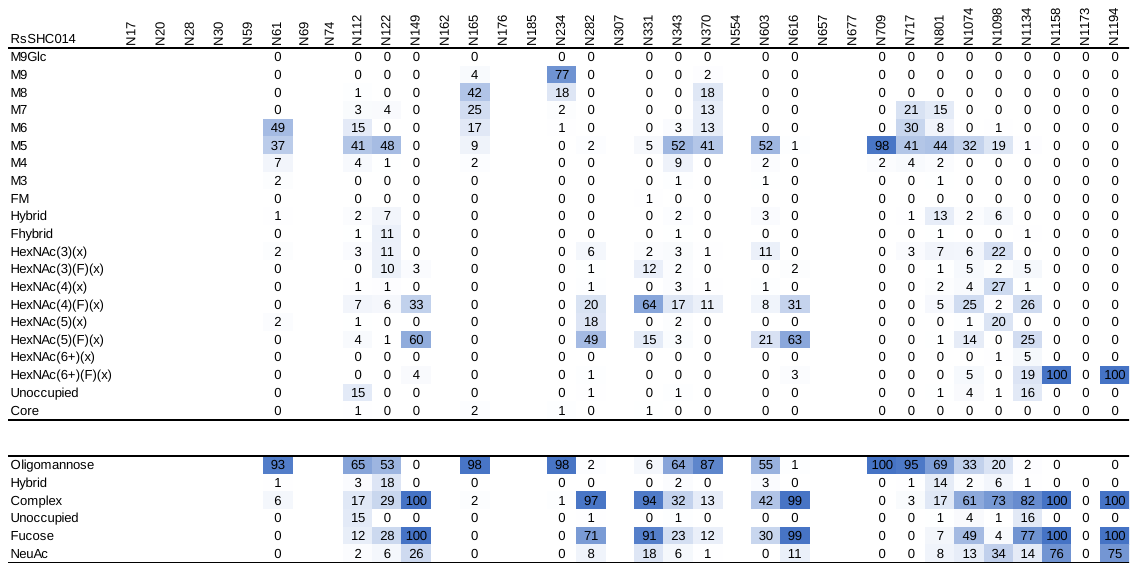
**Supplemental Table 3: Site-specific glycan analysis of clade 1a sarbecoviruses**

**Supplemental Table 4: Site-specific glycan analysis of clade 1b sarbecoviruses**

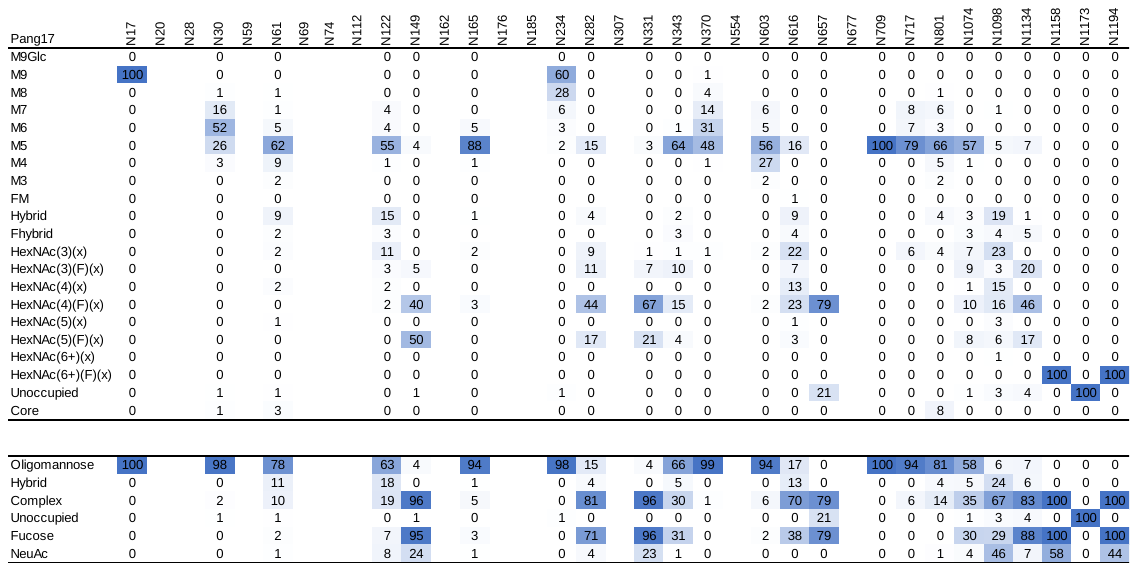

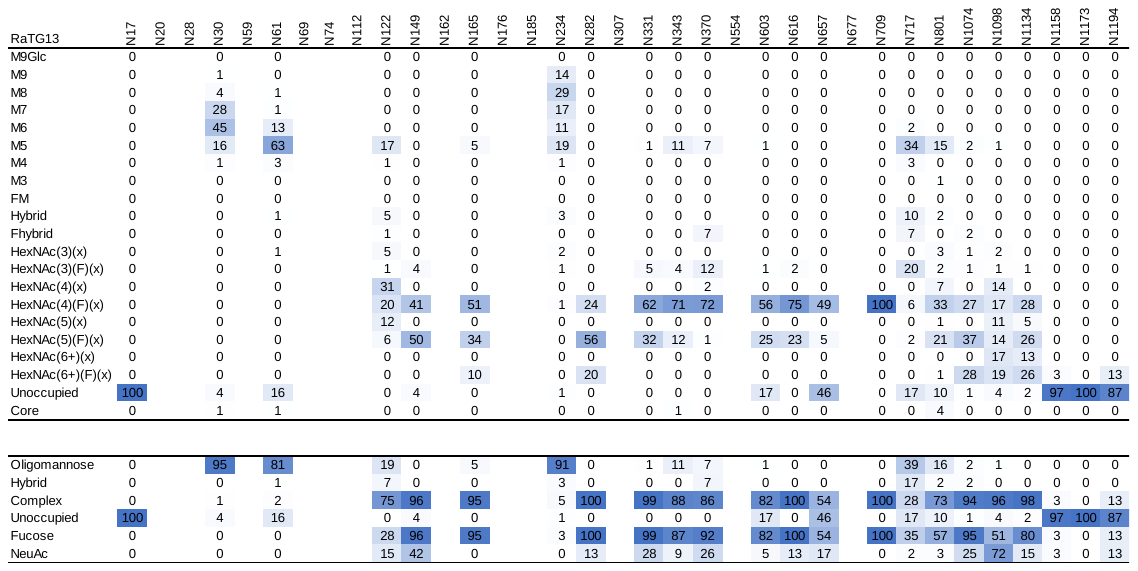

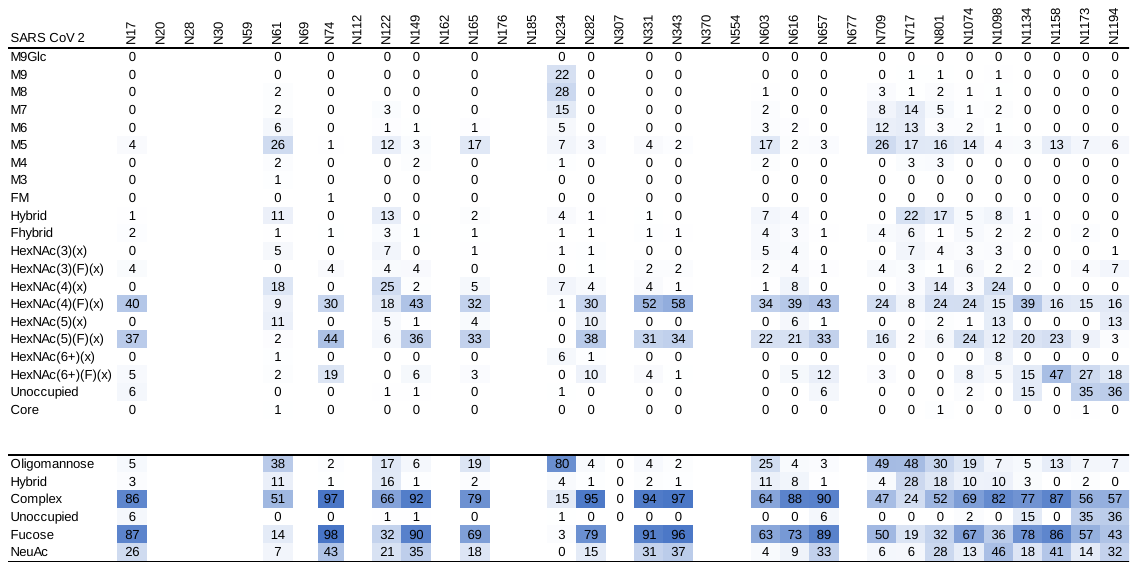

**Supplemental Table 5: Site-specific glycan analysis of clade 2 sarbecoviruses**

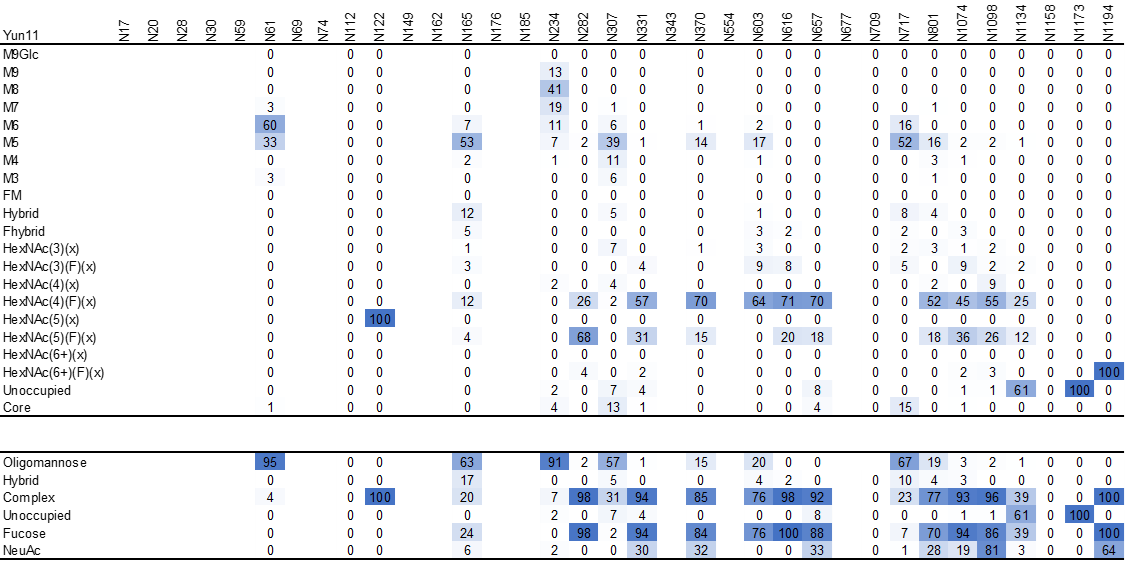

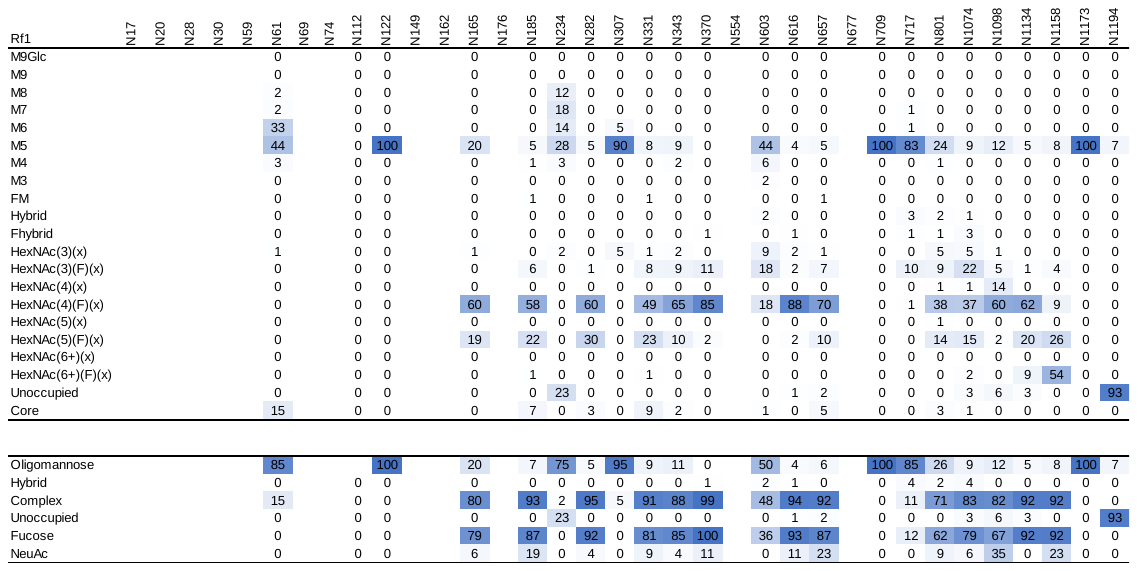

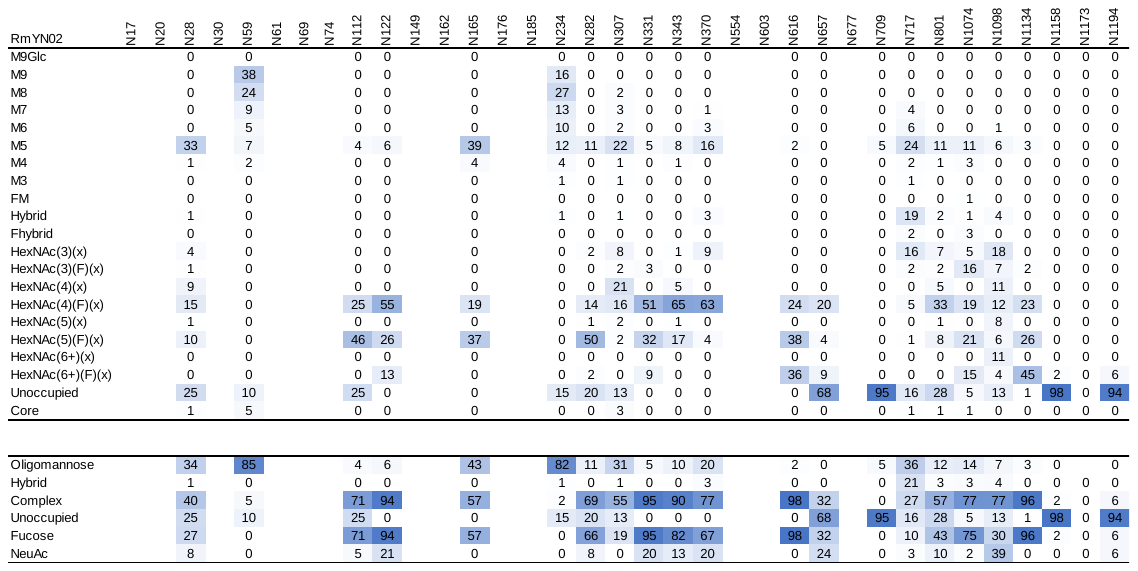

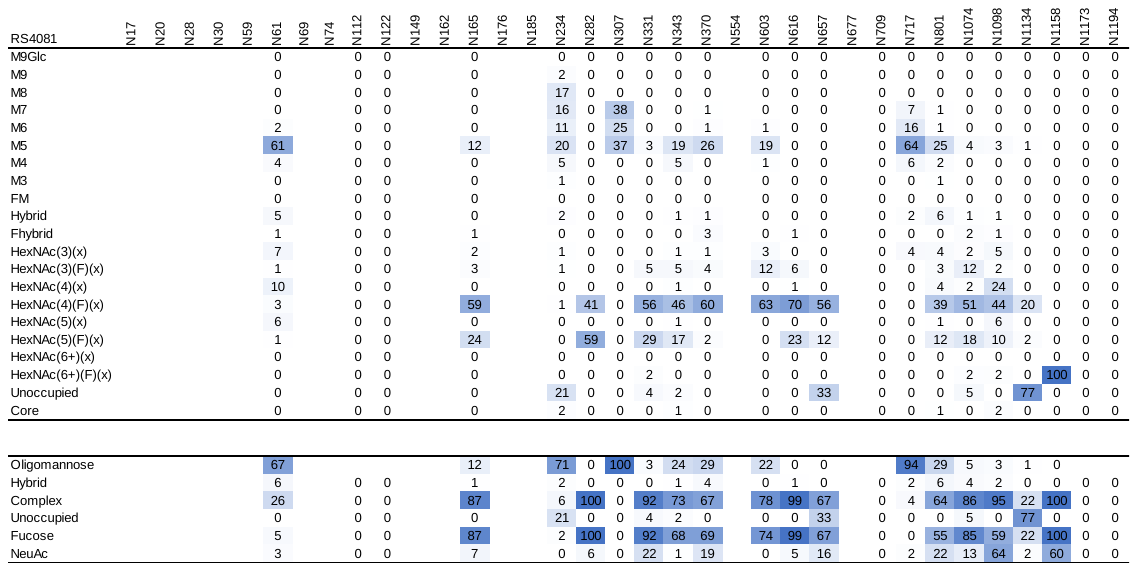

**Supplemental Table 6: Site-specific glycan analysis of clade 3 sarbecoviruses**

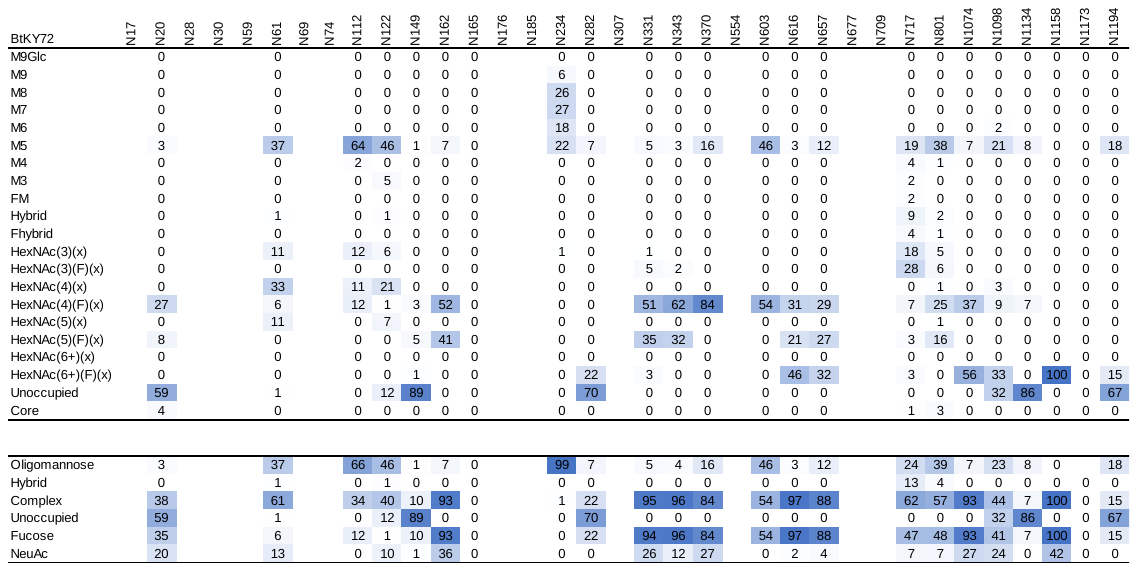

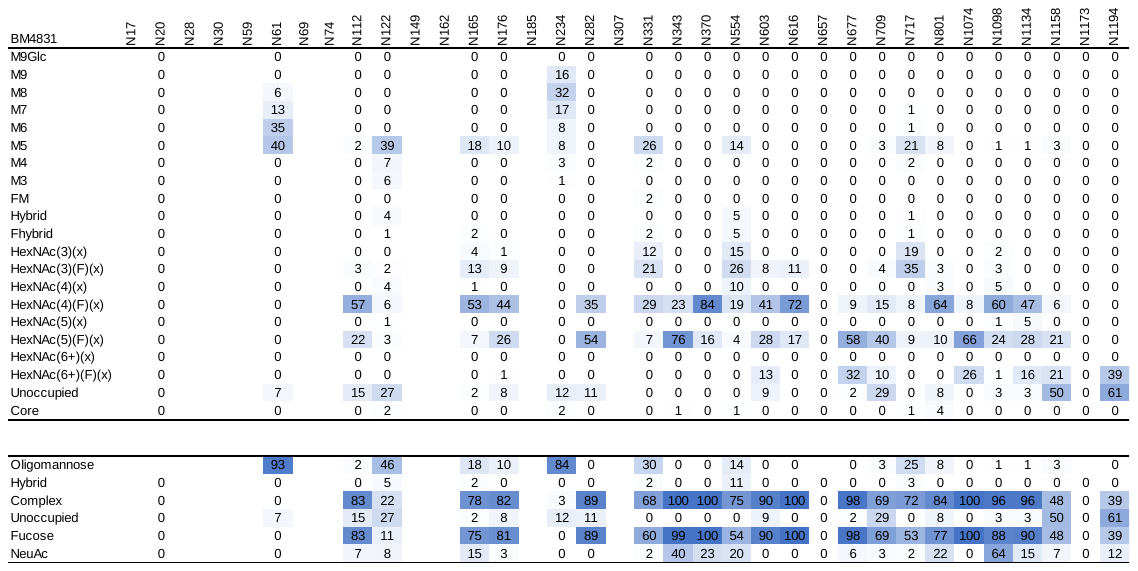
